# AliceDB database and pipeline for identification of natural protein variants based on mass spectrometry measurement data

**DOI:** 10.64898/2026.06.11.731579

**Authors:** Marcel Thiel, Alicja Różycka, Michał Puchalski, Stanisław Ołdziej

**Affiliations:** Intercollegiate Faculty of Biotechnology Gdańsk University and Medical University of Gdańsk, Abrahama 58, 80-307 Gdańsk, Poland

**Keywords:** proteomics, peptidomics, de novo search, natural variants, mutation

## Abstract

The natural variation that distinguishes living organisms within a single species is currently being studied intensively, primarily at the genetic level. Unfortunately, studies of natural variants at the level of protein gene products are not very common, mainly due to the lack of appropriate databases and bioinformatics tools. The main research technique used to study proteomes/peptidomes is mass spectrometry (MS). A classic method for interpreting raw mass spectrometry data in proteomic/peptidomic studies involves the use of databases containing representative (canonical) sequences that define the proteome of the organism under study. In this paper, we present the AliceDB database, which contains information on over 7 million natural variants of protein sequences described in the scientific literature for Homo sapiens. The data contained in the AliceDB database can be utilized using widely available and commonly used software for interpreting proteomic data. Test results regarding the use of the AliceDB database for the interpretation of proteomic data indicate that accounting for the presence of natural variants increases both the number and quality of identified proteins. Furthermore, it is easy to identify protein sequence variants that may, for example, be of significance in medicine.

## Introduction

Genomic population studies indicate that the individual genomes of *Homo sapiens* differ from the reference genome at approximately 4.1–5 million loci. The vast majority of these differences/variants (over 99%) are SNPs (single nucleotide polymorphisms), with a small number of larger changes involving larger segments of genetic material [1]. If we limit ourselves to the part of the genetic material comprising only exonic fragments (so-called coding sequences), the frequency of differences relative to the reference genome is one difference per approximately eight nucleotides [2]. The data presented in the works cited above demonstrate significant variability in genetic material within a population of a single species [1,2], and information on this individual variation in genetic material is increasingly being utilized in medicine, e.g., oncology [3]. While research on the diversity of genetic material within a population is quite common, the study and use of information on diversity at the level of protein gene products is not widespread. In research practice, the primary method used for the identification (including sequencing) and quantitative analysis of proteins is mass spectrometry [4]. An essential component of the process of interpreting data derived from mass spectrometry measurements are amino acid sequence databases, e.g., the UniProt database [5]. Unfortunately, standard amino acid sequence databases (e.g., UniProt) contain only a set of so-called canonical (reference) sequences; as a result, amino acid sequences containing variants are practically ignored in such analyses. Of the many examples in the literature where protein variants are identified in proteomic studies, such research is most often based on amino acid sequence databases created specifically for particular studies using available genomic data [6]. This approach (amino acid sequence databases built ad hoc for specific studies) has many drawbacks, such as cost and the limitation of studies to specific sample types, patient groups, etc. Recently, the first universal database of protein variant sequences, ProHap [7], was launched, based on genetic data from the 1000 Genomes Project [1]. The ProHap database is the first tool that allows for the effective identification of protein variants, but it has significant limitations. The most important one is that the data included in the ProHap database is based solely on the 1000 Genomes Project [1], whereas the available data on protein variants is much more extensive. In this paper, we present a database (AliceDB) based on a much broader resource of information regarding natural variants, as well as a pipeline allowing for the rapid construction of a natural variant database dedicated to specific research.

## Material and Methods

### Data extraction

Protein sequences (both canonical and isoforms) of *Homo sapiens* were obtained from Proteomes collections subdivision of the UniProt database (Release 2025_01) using streaming API endpoint. As a result, 104949 protein sequences were fetched (82861 canonical and 22088 isoforms, respectively). For each UniProtID, EBI Proteins API [8] was used to gather information about protein variants with source filtration to keep those belonging to the ClinVar [9], NCI-TCGA [10], NCI-TCGA Cosmic [3], 1000Genomes [1] and UniProt only. As of that, 7179974 variants were selected for further analysis (see Table 1S). To get additional information about phenotype and pathogenicity of each variant, variants were queried against source databases (e.g., all records related to the ClinVar Databse were queried against its source data gathered from ClinVar FTP server).

### Transformation

All records from data extraction step were cleared and filtered to contain only data needed in further analysis where any kind of metadata associated with records like origin/organism name, variants codon changes etc. were removed. During the transformation process variant types (i.e. mismatch, insertion, deletion, frameshift, duplication etc.) were matched on the corresponding canonical sequences used as templates. Both canonical/isoform and variants were assigned with their own and unique id numbers (called AliceID), where “as” prefix is used to indicate canonical/isoform sequences and “av” to variants.

### Load

After the transform step all records were loaded to the two internal databases. PostgreSQL RDBMS [11] was used to store all data with differentiation for canonical/isoforms and variants (they were stored in separate tables). Elasticsearch [12] was used as a database/search engine where an index was created to accept wildcard queries. To allow scheduling and manage ETL pipeline workflow, Apache Airflow [13] was used.

## Functionality of the AliceDB Database

The AliceDB database is available at www.alicedb.ug.edu.pl. No prior registration is required to use the database, and access is limited only by the performance of the computing resources supporting the database. The database can be searched in several ways. 1) Using a protein sequence identifier from the UniProt database. For such a search, the results include the canonical sequence and, if applicable, a list of isoforms, as well as a list of all natural variants available in the database (see Figure 1S). 2) Using identifiers from the AliceDB database (AliceID), with the search results being basic data, i.e., information on whether it is a canonical sequence, an isoform, or a natural variant (see Figure 2S). 3) Using a peptide sequence; in this case, the search results are one or more records (AliceID) containing the amino acid sequence of the searched peptide (see Figure 3S). In addition to searching for individual peptides, it is possible to search a database using a list of peptides. This option is particularly useful for interpreting data from mass spectrometry measurements. A detailed description of searching the database using a list of peptides can be found in the Supplementary Data (see Figure 4-5S and related text in Supplementary Data). The result of a search using a peptide list is a list of records (AliceIDs) containing the sequences of the searched peptides and a list of complete amino acid sequences (canonical sequences, isoforms, sequences containing natural variants) in FASTA format (see Suplemental Data).

### Pipeline for protein variants identification

The proposed procedure for identifying natural protein variants requires the use of software capable of de novo interpretation (without the use of an amino acid sequence database) of raw data derived from MS measurements. In the examples shown in this paper, the commercial software PeakStudio v. 12.5 Bioinformatics Solutions Inc. Waterloo, Canada [14-15] was used. However, any software capable of de novo peptide identification can be used in proposed approach [16].

Figure 1 presents pipeline for protein variants identification proposed in this work. Raw data from MS measurement were initially analyzed using Peaks Studio ver. 12.5 software [16] with using *Homo Sapiens* protein sequences from the UniProt database as a target sequences (see Supplemental Data for details). Next, peptides sequences both identified based on database search, and de novo sequences were merged to form one input. Combined set of peptides (canonical+ de novo) were searched against data from AliceDB (peptide list search option). As results of AliceDB search custom made database of sequences (canonical as well those containing natural variants) is created. The new custom-made database of sequences is used by Peaks Studio to second round of analysis of raw input data.

**Figure 1.**
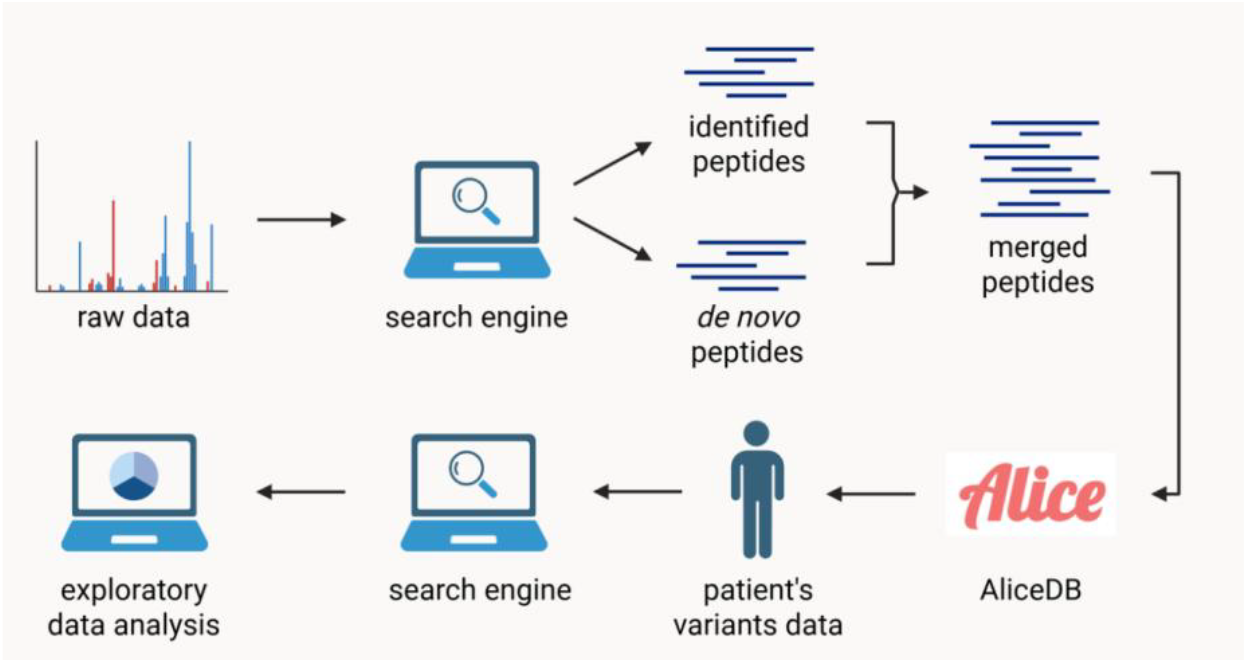
pipeline for protein variants identification proposed in this work. See accompanying text for details.

## Results

To tests proposed database and data processing pipeline data retrieved from the PRIDE repository [17] record PXD024347 were used [18]. Detailed information regarding raw mass spectra data processing are provide in the Supplementary Data.

Table 1 presents numerical data comparing the annotation of raw MS measurement data using the UniProt database with annotation using the AliceDB database and UniProt database. It can be observed that thanks to the use of the proposed solutions (AliceDB), the overall number of protein sequence identifications increases by approximately 20–25% compared to classical methods using the UniProt database (the last two (6–7) rows in Table 1). This significant increase in identifications is due to two factors. First, the identification of sequences containing variants (better sequence coverage, a greater number of identified peptides assigned to a given protein sequence), but also the fact that the AliceDB database contains isoform sequences, whereas the version of the UniProt database used in this study contains only canonical sequences (typically, this version of the UniProt database is used in proteomic studies). Nevertheless, it is worth noting that out of the total pool of peptides identified in the samples (first row of Table 1), approximately 5% of the peptides can be uniquely assigned only as fragments of proteins that are natural variants of canonical sequences (see row 5 of Table 1). These approximately 5% of peptides (row 5, Table 1) would have been omitted using classical searches with standard methods. Furthermore, approximately 3–5% of the peptides identified in the sample, even when using the AliceDB database, are not annotated (the difference between the number of peptides in rows 1 and 2 of Table 1). This still unidentified pool of peptides is likely mostly due to the incompleteness of available information on natural variants, which highlights further opportunities for the development of the presented project.

**Table 1.** Summary and comparison of raw MS data annotations using the standard procedure based on the UniProt database, and the pipeline proposed in this paper.

| Samples<sup>1</sup> | 114 | 123 | 125 | 126 | 119 | 127 | 128 | 131 |
| --- | --- | --- | --- | --- | --- | --- | --- | --- |
| 1.Number of peptides identified using UniProt database + peptides identified by de novo search | 25247 | 26080 | 26402 | 27860 | 26488 | 25591 | 24442 | 25066 |
| 2. Number of peptides annotated in AliceDB database | 23940 | 25260 | 25753 | 27060 | 25292 | 24691 | 23260 | 23675 |
| 3.Number of peptides with possible variants | 6722 | 6722 | 5732 | 6195 | 6450 | 6291 | 6435 | 6370 |
| 4. Number of unique peptide with variants<sup>2</sup> | 1333 | 1286 | 1051 | 1150 | 1306 | 1208 | 1324 | 1289 |
| 5. Number of proteins identified based on UniProt database | 1643 | 3225 | 2164 | 2620 | 2647 | 1739 | 1802 | 1779 |
| 6. Number of proteins identified based on AliceDB database | 2031 | 3662 | 2291 | 2823 | 3175 | 2161 | 2228 | 2284 |

<sup>1</sup>Samples are marked according to symbols used in record PXD024347 from PRIDE repository [18].
<sup>2</sup>Peptides containing sequences that are potentially those of natural variants, provided that peptides shorter than 9 amino acid residues are excluded, as well as peptides whose sequences are identical to canonical sequences other than those of the protein in question.

The examples in Table 2 show that taking variants into account increases the number of identified peptides that can be considered for the identification of a given protein sequence. An increase in the number of peptides assigned to a given sequence results in a higher coverage rate (approximately 2–6%). The examples also show that the number of identified peptides containing a variant is usually greater than one, which means that the identification of the variant itself is quite reliable. Peptides of different lengths cover the same sequence region containing the variant. An extreme example of variant inclusion is the P06315 protein. Using only the UniProt database identifies only one peptide, which means that the reliability of the protein identification does not meet the quality and statistical criteria [19]. However, using the AliceDB database allows for the identification of an additional peptide in a different region of the sequence, thereby meeting the identification criteria. An interesting example is protein P02774. According to data from the NCBI database [20], the canonical sequence deposited in the UniProt database occurs in only about 0.2% of the population, whereas the identified variant is dominant and occurs in about 99.8% of the population.

**Table 2.** Examples of identified natural variants in the analyzed samples.

| UniProtID | Peptides/<br>Coverage<sup>1</sup> | AliceID | Peptides/<br>Coverage<sup>2</sup> | Variant<sup>3</sup> |
| --- | --- | --- | --- | --- |
| P02774 | 123/88,82 | av0343186 | 128/90,51 | H445R |
| P06315 | 1/10,36<sup>4</sup> | av2487130 | 2/16,52 | T94R |
| P06727 | 71/88,89 | av0372344 | 76/90,15 | S147N |
| P22792 | 49/53,58 | av0554845 | 51/55,23 | Q509R |

<sup>1</sup>Number of peptides identified using the UniProt database / sequence coverage in percent
<sup>2</sup>Number of peptides identified using the AliceDB database / sequence coverage in percent
<sup>3</sup>Observed sequence change in the identified variant
<sup>4</sup>No protein identification based on a single peptide

## Conclusion

The presented database of natural protein variants (AliceDB) contains over 7 million records and is currently the largest collection of its kind, though it is likely still far from providing a complete description of biological diversity of human proteome. The data shown in Table 1 clearly indicate that there remains a pool of peptides that have not been described, which points to the need for further development of both the database and its associated tools. The notation used to describe variants is based entirely on canonical sequences and the identifiers of these sequences from the UniProt database. This distinguishes it from the ProHap database [7], in which the description of protein gene products is based entirely on genetic data. Basing the description of protein variants on notations from the UniProt database significantly facilitates the interpretation of results and the use of classical data processing tools employed in proteomics/peptidomics studies. The adopted method of raw data processing requires two-step processing (see Figure 1 and related text). However, this appears to be a good compromise compared to the alternative of processing data using a sequence database exceeding 7 million entries, which can be cumbersome given users’ limited hardware resources. The ability to create personalized databases for a single sample or group of samples, as proposed in this work, appears to be an interesting option when searching for specific protein variants in numerous samples (e.g., in medical research). Currently, the AliceDB database offers data only for the species *Homo sapiens*, but the IT solutions employed allow for the database to be expanded to include any other species.

## Authors’ contribution

MT, AR developed database, MT, MP and SO preformed analyses and drafted the paper. All authors read and approved the final version of manuscript.

## Acknowledgements

The research was partially funded by the University of Gdańsk’s Small Grants Program—UGrants-start 5 (MP). Computational resources used in this project were provided by the Informatics Center of the Metropolitan Academic Network (IC MAN-TASK) in Gdańsk.

