## Supplemental_data for "AliceDB database and pipeline for identification of natural protein variants based on mass spectrometry measurement data"

### Supplemetal data

**Table 1S.** The number of natural variants stored in the current version of AliceDB database, with variants classified by the processes occurring in the genetic material.

| Variant type | Number of variants in database |
|---|---|
| Missense | 6731831 |
| Stop gained | 340985 |
| Frameshift | 96101 |
| Stop lost | 5548 |
| Inframe deletion | 4536 |
| Insertion | 1762 |
| Delins | 169 |
| Duplication | 85 |
| Repeat | 36 |

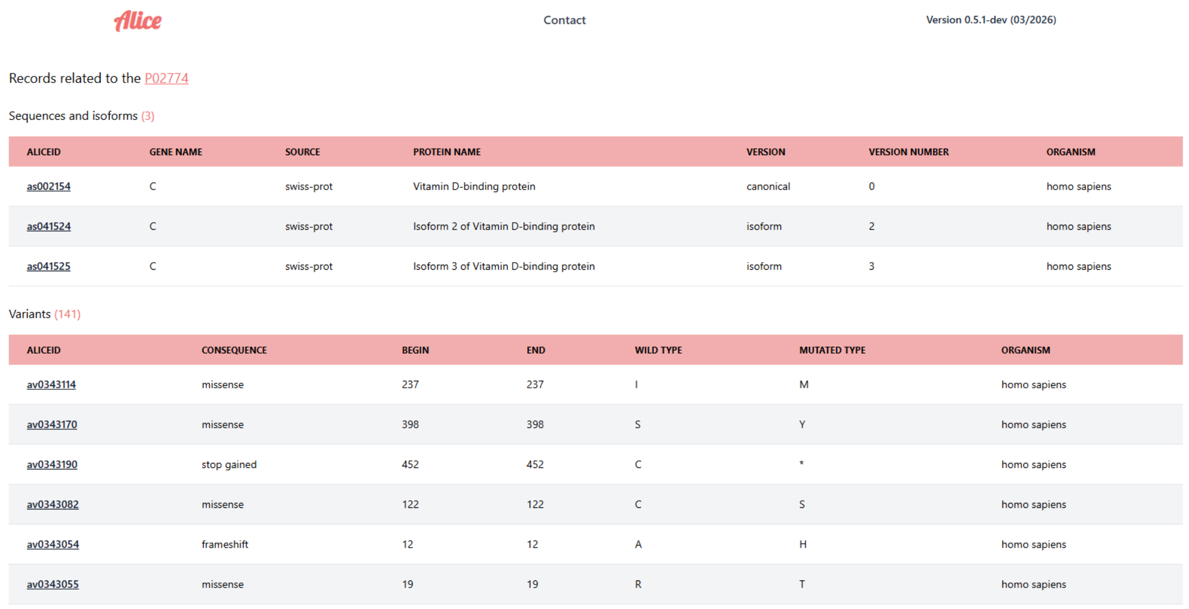

Alice | Contact | Version 0.5.1-dev (03/2026)

Records related to the P02774

Sequences and isoforms (3)

| ALICEID | GENE NAME | SOURCE | PROTEIN NAME | VERSION | VERSION NUMBER | ORGANISM |
|---|---|---|---|---|---|---|
| as002154 | C | swiss-prot | Vitamin D-binding protein | canonical | 0 | homo sapiens |
| as041524 | C | swiss-prot | Isoform 2 of Vitamin D-binding protein | isoform | 2 | homo sapiens |
| as041525 | C | swiss-prot | Isoform 3 of Vitamin D-binding protein | isoform | 3 | homo sapiens |

Variants (141)

| ALICEID | CONSEQUENCE | BEGIN | END | WILD TYPE | MUTATED TYPE | ORGANISM |
|---|---|---|---|---|---|---|
| av0343114 | missense | 237 | 237 | I | M | homo sapiens |
| av0343170 | missense | 398 | 398 | S | Y | homo sapiens |
| av0343190 | stop gained | 452 | 452 | C | * | homo sapiens |
| av0343082 | missense | 122 | 122 | C | S | homo sapiens |
| av0343054 | frameshift | 12 | 12 | A | H | homo sapiens |
| av0343055 | missense | 19 | 19 | R | T | homo sapiens |

**Figure 1S.** A screenshot showing the search results of the AliceDB database using an identifier from the UniProt database (the example uses identifier P02774). The search results include the canonical sequence and isoform sequences (if described) as well as the number and list of known variants (the example shown displays only a portion of the list of 141 variants deposited in the database). Each sequence (canonical, isoform, or natural variant) in the AliceDB database is described using a unique identifier. The identifier consists of two letters and seven digits. The first two letters of the identifier indicate the record's content: whether it refers to a canonical sequence or isoform (prefix "as") or to a natural variant (prefix "av").

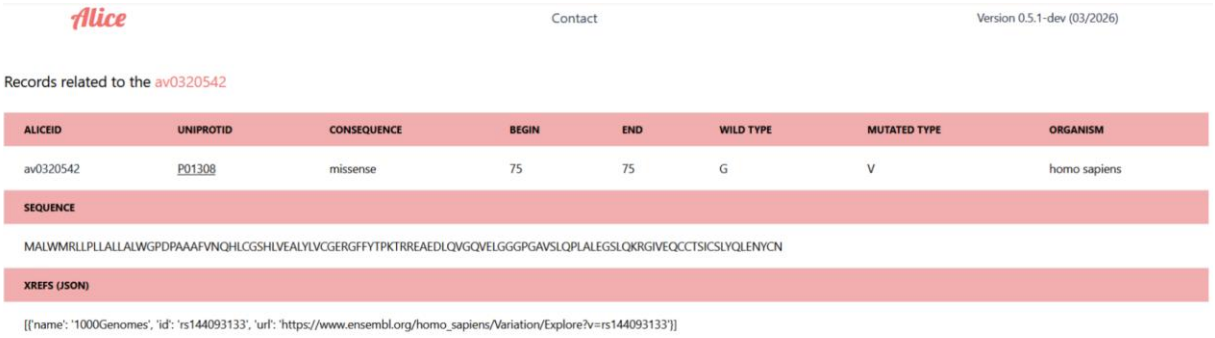


**Figure 2S.** A screenshot showing the results of a search in the AliceDB database using an identifier from the AliceDB database (the example uses the identifier av0320542). In the example provided, a record describing a natural variant was searched for (prefix “av”; see description in Figure 1S). The return information includes a description of the sequence, the amino acid sequence, and a link to literature or databases describing the searched variant.

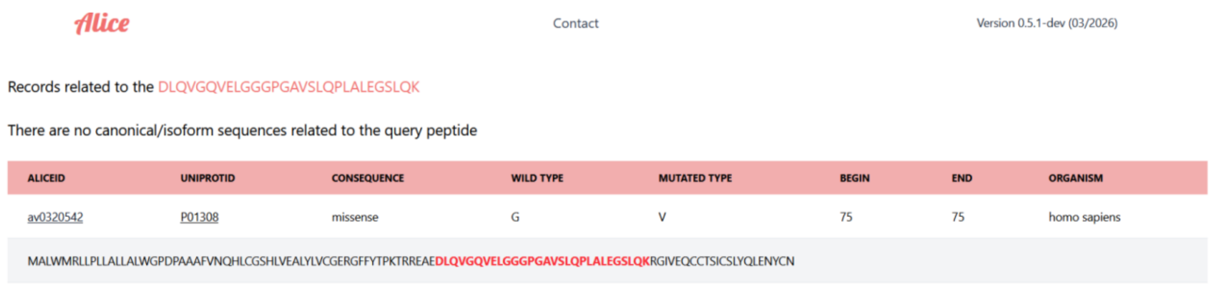


**Figure 3S.** A screenshot showing the search results from the AliceDB database for a peptide sequence. The result of such a search is a sequence or a list of sequences that contain the searched-for peptide.

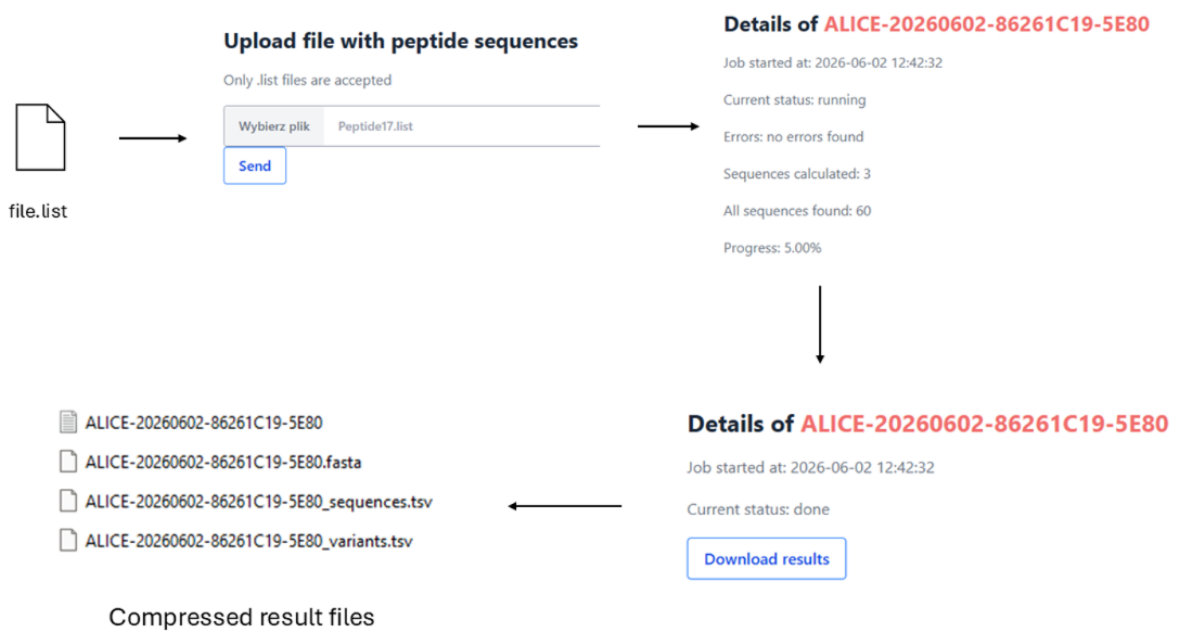


**Figure 4S.** Flowchart of the search process using a file containing a list of peptides. The search data consists of a list of peptides (list.file); after it is loaded into the search engine database and the search is initiated, the task is assigned a unique identifier (shown in red in the figure above). This identifier allows users to track the progress of the search (from the database's main menu, select the "Alice Job" option and enter the search identifier). Once the search is complete, it is possible to download the search results. The search results are returned as a compressed file (zip) containing four files, the contents of which are described below.

Five files are included with this manuscript as a search example in compressed file named Additional_Data.zip:

Peptide17.list - a text file containing sample input data, peptide list

ALICE-20260602-86261C19-5E80 - a file containing the search information

ALICE-20260602-86261C19-5E80.fasta – a file containing sequences in FASTA format for all proteins identified based on input data. This file will provided customised database of protein sequences for given sample. File will includes sequences of all canonical, isoform and natural variants sequences identified in searched peptide list.

ALICE-20260602-86261C19-5E80_sequences.tsv – a file (in tsv format) containing information about identified canonical sequences and isoforms. Every record in this file includes the following information:

QuerySequence - the peptide sequence used to search the database

AliceID - the sequence identifier in the AliceDB database

UniprotID - identifier from the UniProt database

Gene - name of the gene encoding the protein

Source - location of the sequence in the Swiss-Prot or Trembl database

ProteinName - name of the protein

VersionNumber – in the UniProt database, some sequences that are isoforms have their own identifiers, but some isoforms are designated as, for example, Q71UI9-2, Q71UI9-3. If this parameter is equal to 2 or more, it indicates additional isoforms assigned to a given UniProt record. A value of 0 indicates the canonical sequence

Version - indicates whether we are dealing with a canonical sequence or an isoform

Positions - the location of the peptide (QuerySequence) within the protein sequence

Organism - currently, the database is for the Homo sapiens organism only

Sequence - the entire protein sequence

ALICE-20260602-86261C19-5E80_variants.tsv – file (in tsv format) containing information about identified natural variant sequences. Every record in this file includes the following information:

QuerySequence - the peptide sequence used to search the database

AliceID - the sequence identifier in the AliceDB database

UniprotID - the identifier from the UniProt database

WildType - the amino acid residues found at a specific position in the canonical sequence.

MutatedType - amino acid residues found in the natural variant

Begin End - describe the range of residues in the canonical sequence affected by the variation

Consequence - variant type; see Table 1S.

Organism - currently, the database is operational for the Homo sapiens organism only

Sequence - the entire amino acid sequence containing the variant

**Raw mass spectra processing**

Raw results data from MS measurement were downloaded from PRIDE repository PXD024347 [1] and analyzed using Peaks Studio 12.5 [2] with using *Homo Sapiens* protein sequences from the UniProt database (Swiss-Prot + TrEMBL accessed 16.01.2025) as a target sequences in Spider Search workflow. Precursor mass error tolerance was set for10 ppm and fragment mass error tolerance was 0.02 Da. Digest mode was semi-specific, with enzyme specified by sample (trypsin, chymotrypsin or Lys-C) and maximum 2 missed cleavages. Peptide length range was set for 6-45 amino acids. Carbamidomethylation, carbamylation, deamidation (NQ), dormylation, oxidation (M) and pyro-glu from Q were chosen as variable modifications, with maximum 2 modifications per peptide. For De Novo peptide identification parameter ALC was set for 15%. Peptides sequences identified in database search and de novo only search were merged to form one input list analyzed by AliceDB (see Figure 1 in main text). After that, all variants identified by AliceDB search were used to extend the database used by Peaks Studio (fasta sequences of proteins with identified variants were added to previously used database) and raw input was analyzed once again (against new/extended database) with the same search parameters as in first pass of database search.

References.

1. Lewandowska, A.E.; Fel, A.; Thiel, M.; Czaplewska, P.; Łukaszuk, K.; Wiśniewski, J.R.; Ołdziej, S. Compatibility of Distinct Label-Free Proteomic Workflows in Absolute Quantification of Proteins Linked to the Oocyte Quality in Human Follicular Fluid. *Int. J. Mol. Sci.* **22**, 7415. (2021) https://doi.org/10.3390/ijms22147415

2. Zhang J., Xin L., Shan B., Chen W., Xie M., Yuen D., Zhang W., Zhang Z., Lajoie G.A., Ma B. PEAKS DB: De novo sequencing assisted database search for sensitive and accurate peptide identification. *Mol. Cell. Proteom.* **11**, M111.010587 (2012) https://doi: 10.1074/mcp.M111.010587
